# Mechanical and elemental characterization of ant mandibles: consequences for bite mechanics

**DOI:** 10.1101/2023.09.20.558673

**Authors:** Cristian L. Klunk, Michael Heethoff, Jörg U. Hammel, Stanislav N. Gorb, Wencke Krings

## Abstract

Chewing with the mandibles is a food processing behavior observed in most current insect lineages. Mandible morphology has an essential role in biting behavior and food processing capacity. However, the mandible cuticle can have regional differences in its mechanical properties, associated or not with the accumulation of elements that increase cuticle stiffness. The effects of such a heterogeneous distribution of cuticle material properties in the mandible responses to biting loading are still poorly explored in chewing insects. Here we measured the elemental composition and material properties of workers of an ant species, *Formica cunicularia*, and tested the effects of the cuticular variation in Young’s modulus (E) under bite-loading with Finite Element Analysis (FEA). We divided worker mandibles into four regions that we expect would vary in elemental composition and material properties, namely the masticatory margin, mandible blade, ventral (VMA), and dorsal (DMA) mandibular articulations with the head. Specifically, we expect the masticatory margin will show higher cuticular hardness (H) and E values, followed by the mandibular joints and the mandible blade. We also predict that such cuticle material properties variation is functionally relevant under bite-loading, changing stress patterns when compared to the mechanical responses of a mandible with a homogeneous distribution of material properties. To measure elemental composition, we used energy disperse X-ray spectroscopy, while H and E were accessed through nanoindentation tests. Mandible mechanical responses to bite-loading were tested with FEA, comparing a mandible with a homogeneous versus a heterogeneous E distribution. As expected, the mandibular regions showed distinct proportions of relevant elements, like Cu and Zn, with the masticatory margin showing the higher levels of those elements, followed by the mandibular articulations with the head and the mandible blade. The same pattern was observed regarding the values of cuticle H and E. When incorporated into FEA, this variation in E effectively changed mandible stress patterns, leading to a higher concentration of stresses in the stiffer mandibular regions, letting the softer mandible blade with relatively lower stress levels. Our results demonstrated the relevance of cuticle heterogeneity in mechanical properties to deal with bite-loading demands and suggest that the accumulation of transition metals such as Cu and Zn has a relevant correlation with such mechanical characteristics of the mandible in this ant species.

## Introduction

Many insects evolved the ability to capture prey and chew food items using their mandibles (Snodgrass 1935, Blanke 2019). However, different species also employ their mandibles to perform other tasks besides food processing, like intraspecific contests, defense, transportation, construction, and excavation. Such a multitask use of the mandibles is commonly observed among ant workers (Wilson 1987; Zhang et al. 2020; Richter & Economo 2023), responsible for most of the colony’s non-reproductive activities. Ants show a relevant interspecific variation in mandible morphology (Sosiak & Barden 2021), whereas intraspecific distinctions are also observed, mainly between worker types (Casadei-Ferreira et al. 2021). How morphological disparity reflects ant biting performance was investigated for some ant lineages by the application of computational simulations (Larabee et al. 2018; Zhang et al. 2020; Klunk et al. 2021; Wang et al. 2022), as well as through the estimation of relevant mechanical characteristics based on morphological information (Püffel et al. 2021; 2023a), providing compelling evidence about how mandibular morphological variation can influence bite mechanics.

An approach widely employed to investigate form-function relationships in biological structures is Finite Element Analysis (FEA). It consists in a numerical method that approximates the structure’s mechanical responses to external loads, revealing patterns of structure deformation, strain, and stress (Rayfield 2007). A digital representation of the target structure, knowledge about its material properties, and information on its boundary conditions (regions of loading and contact with other structures) are necessary to define an FEA (Rayfield 2007). Regarding insect mandibles, FEA was employed to investigate its mechanical behavior in Odonata (Blanke et al. 2017b), beetles (Goyens et al. 2015), and ants (Larabee et al. 2018; Zhang et al. 2020; Klunk et al. 2021; Wang et al. 2022) considering different behaviors related to biting. However, in those cases, the mandible cuticle was modeled as a homogeneous material, even though it is recognized that the mechanical properties of insect cuticles can vary substantially along the same structure (Schofield et al. 2002; 2021; Vincent and Wegst, 2004; Brito et al. 2017; Li et al. 2020; 2022a; 2022b; Ma et al. 2020), with relevant biomechanical effects (Schofield et al. 2002; 2021; Rajabi et al. 2017; Das et al. 2018; Jafarpour et al. 2020; Li et al. 2020; Matsumura et al. 2020; Casey et al. 2022). In an FEA case study on mollusks teeth, which were either homogeneous or heterogeneous in material properties, the effect of the mechanical property distribution on teeth stress and strain patterns could be documented (Krings et al. 2020). It showed that when the material was heterogeneous, stresses increased and strains decreased, whereas teeth of homogeneous material experienced less stress levels and higher strains.

In comparative studies of animal biomechanics using FEA, it is usual to assume that the structure material properties are homogeneously distributed (Marcé-Nogué 2022) and that its interspecific variation is negligible or is only not the focus of the study. This assumption is not a problem only when the intention is to perform a comparative analysis on objects varying little in their material properties and to investigate the sole effects of morphological variation (Rayfield 2007). However, an ideal approach in organismal biomechanics and functional ecology should explore the role of material properties variation, which would also aid in our understanding of the interplay between materials and structure and provide some insight into the material properties’ evolution (Rajabi et al. 2017; Das et al. 2018; Jafarpour et al. 2020; Li et al. 2020; Matsumura et al. 2020; Casey et al. 2022; Krings et al., 2022a). However, researchers usually stumble on the scarcity of data regarding the mechanical properties of biological materials, a consequence of the difficulties associated with such measurements, and the impressive intraspecific and even intraindividual variation that reduces the representativeness of such data at the species level (Stamm et al. 2021).

Two main components of the insect cuticle are chitin, a polysaccharide polymer, and a series of proteins (Zhu et al. 2016). One of the most relevant aspects of the cuticle is its sclerotization or tannification, which involves chemical reactions in the exocuticle that change the molecular arrangements between chitin and the protein matrix, modifying the mechanical properties of this cuticular layer, like its stiffness, hardness, breaking stress, among other characteristics (Vincent & Wegst 2004; Neville 2012). In addition, transition (Cu, Fe, Mn, Zn) and alkaline earth metals (Ca, Mg) can bind strongly to the polymers increasing the cross-linking density (Vincent 2002; Lichtenegger et al. 2003; Waite et al. 2004; Quicke et al. 2004; Pontin et al. 2007; Broomell et al. 2008; Degtyar et al. 2014; Liu et al. 2017; Polidori & Wurdack 2019).

Insect cuticle has essential structural and protective functions, providing the necessary support for muscle anchoring. Also, it protects mechanically and chemically the insect from the environment (Schroeder et al. 2018; Buxton et al. 2021). Several insect behaviors impose relevant mechanical demands on their exoskeleton, like flying, jumping, running, walking, biting, and the associated muscle contractions. Such behaviors usually generate friction between body parts and the environment and even among body parts that can lead to cuticular wear, which could also happen with the frequent use of a structure like the mandibles to process hard materials (Schofield et al. 2011; Nadein and Gorb, 2021; 2022; Nadein et al., 2021; Püffel et al. 2023b). Therefore, it is not surprising that substantial variation in cuticle material sclerotization levels is observed along the body of an insect (Michels & Gorb 2012; Büsse & Gorb 2018; Eshghi et al. 2018; Li et al. 2020; Josten et al. 2022; Krings & Gorb, 2023), besides the differences among the cuticular layers or the abundance of transition or alkaline earth metals. This intraindividual variation in the cuticle can have significant functional relevance due to its effects on the cuticle material properties (Das et al. 2018; Jafarpour et al. 2020; Li et al. 2020; Matsumura et al., 2020; Toofani et al. 2020; Xing & Yang 2020; Casey et al. 2022).

Among several mechanical parameters that characterize a material, hardness (H) and Young’s modulus (E) provide relevant information on its behavior under loading (Arzt et al. 2002; Enders et al. 2004; Barbakadse et al. 2006; Labonte et al. 2017). Young’s modulus measures the material resistance to elastic deformation under compressive or tensile forces, while H reflects the material resistance to plastic deformation (Berthaume 2016). Both material properties in biological systems can vary substantially intra- and interspecifically and be modified by the degree of sclerotization or the concentration of inorganic elements. Elements like Zn, Mn, Ca, and Mg increase cuticle H and improve the wear resistance of insect appendices (Hillerton et al. 1982; Hillerton & Vincent 1982; Edwards et al. 1993; Schofield et al. 2002, 2021; Cribb et al. 2008; Andersen 2010; Vega et al. 2017; Zhang et al. 2019; Kundanati et al. 2020; Johnston et al. 2022; Krings et al. 2022d). Transition and alkaline earth metals in the cuticle are observed in several arthropod lineages, especially on appendages associated with biting or puncture, like spider fangs (Politi et al. 2021; Schofield et al. 2021), scorpion claws, pedipalps, and chelicerae (Schofield et al. 2003; Schofield et al. 2021), mandibles of termites (Cribb et al. 2007), cicadas (Reiter et al. 2023), ant lions (Krings & Gorb 2023), and ants (Schofield et al. 2002; 2003; 2021; Polidori et al. 2020). Along with these cross-links, it is known that ant species can incorporate minerals into their cuticle (Li et al. 2020), similar to the cuticles of crustaceans. All of this highlights the relevance of determining the cuticle elemental composition, as is being explored in the chitinous structures of several mollusk lineages (Krings et al. 2022a; 2022b; 2022c; 2022e), especially regarding structures heavily employed to perform multiple tasks, like the ant mandibles.

The present study aims to provide a mechanical and elemental characterization of *Formica cunicularia* Latreille, 1798, worker mandibles, and test through FEA how the mandibular variation in cuticle E influences its responses to bite loading. Based on the distribution of stress observed in ant mandibles in previous attempts to simulate biting behavior with FEA (Larabee et al. 2018; Zhang et al. 2020; Klunk et al. 2021; Wang et al. 2022), we divided the mandible into four regions, namely the masticatory margin, dorsal (*DMA*) and ventral (*VMA*) articulations with the head, and the mandible blade. We hypothesize that the masticatory margin will show the highest value of E and H, followed by the mandibular articulations with the head and the mandible blade. Also, we predict this ranking regarding the proportion of transition and alkaline earth metals along the mandible cuticle. Finally, we anticipate that when considering the measured E values in FEA simulations, there will be a difference in relative stress distribution compared to simulations with a homogeneous E definition.

## Material and methods

### Ant specimens

*Formica cunicularia* is a common European ant, especially abundant in Central Europe (Guenárd et al. 2017) and frequently found in urbanized areas. Specimens of *F. cunicularia* were collected from a colony located in an urban area in Darmstadt, Germany, in 2022 and stored in 70% EtOH. They are now deposited at the Leibniz Institute for the Analysis of Biodiversity Change, Leibniz, Germany. It is recognized that the conditions of sampling storage affect the material property values, mainly due to its effects on the sample’s hydration state (Aberle et al. 2017; Li et al. 2022a), but the analyzed specimens were stored under the same conditions and the chemical or mechanical characteristics are thus comparable. We recognize that the specific conditions of those samples’ storage prevent the direct comparison of material properties values with measurements from other efforts available in the literature, but should not prevent us from testing our hypothesis related to the distinct mandibular regions.

### Mandible scans

With appropriate imaging techniques, it is possible to observe gradients of material density that suggest the accumulation of heavier atoms, like heavy metals. Phase contrast imaging methods allow for the detection of heterogeneity in material density, which points to regions with potential differences in elemental composition (deposition of heavy metals) and consequent variation in material properties (e.g. E and H). Mandibles from one *F. cunicularia* worker was scanned using synchrotron radiation X-ray tomography (SRμCT) at the Imaging Beamline P05 (IBL) (Greving et al., 2014; Haibel et al., 2010; Wilde et al., 2016) operated by the Helmholtz-Zentrum-Geesthacht at the storage ring PETRA III (Deutsches Elektronen Synchrotron – DESY, Hamburg, Germany), through a phase contrast method. Tomographic reconstruction was done with a custom reconstruction pipeline (Moosmann et al. 2014) using MatLab (Math-Works) and the Astra Toolbox (van Aarle et al. 2015, van Aarle et al. 2016, Palenstijn et al. 2011). To visualize the distribution of cuticle density and pre-segment the mandible for biomechanical simulations, we imported the scan data to the software Amira 5.4 (Visage Imaging GmbH, Berlin, Germany). Snapshots of mandible cross-sections were taken to illustrate the occurrence of mandibular regions with higher density (whitish voxels). To generate a surface model of the *F. cunicularia* worker mandible we imported the scan data into Amira 5.4 and manually segmented it using the Magic Wand tool at intervals of 10 slices. This pre-segmented mandible file was uploaded to the online platform Biomedisa (Lösel et al. 2020) for automatic interpolation between the pre-segmented slices and the generation of a mandible representation. Finally, the outputs from Biomedisa were imported back into Amira 5.4 to correct for inaccuracies and reduce the complexity of the reconstructed morphology.

### Exocuticle elemental composition

Higher levels of cuticular H and E are usually associated with an accumulation of heavier elements (Brito et al. 2017; Schofield et al. 2021). To investigate the elemental composition of the mandible cuticle, we employed energy disperse X-ray spectroscopy (EDX, EDS) on three *F. cunicularia* workers (six mandibles). Here, we used the one specimen that was previously scanned and two additional ones. To test our main hypotheses that mandibular regions vary in their chemical composition and consequently in their mechanical characteristics, we divided each mandible into four regions, namely the masticatory margin, mandible blade, ventral (VMA), and dorsal (DMA) mandibular articulation with the head (Fig.1).

**Fig.1.**
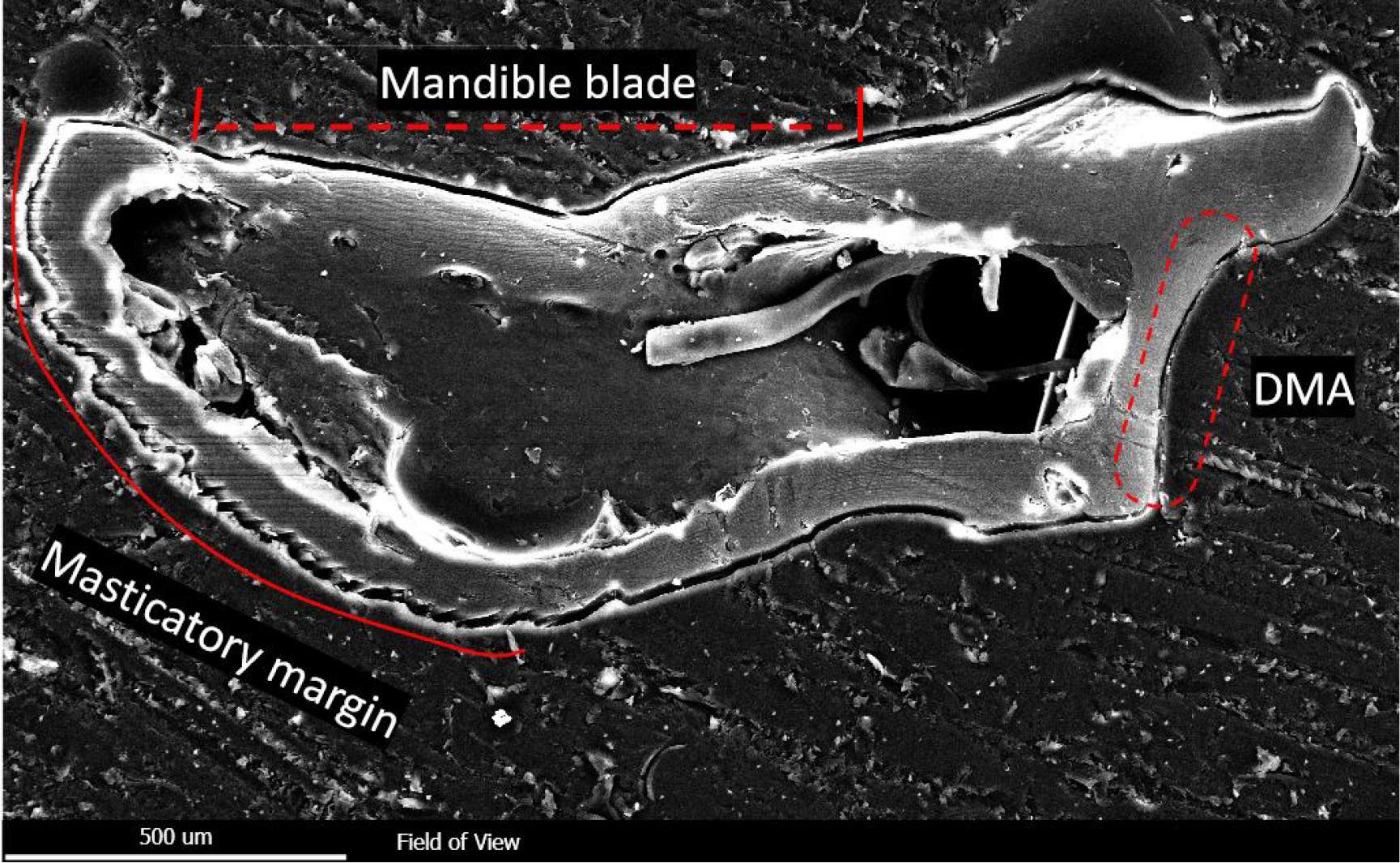
Cross-section of one *F. cunicularia* worker mandible used for the nanoindentation and EDS experiments, highlighting the distinct mandibular regions considered. The ventral mandibular articulation is not visible in this view. DMA = Dorsal mandibular articulation.

First, mandible samples were cleaned in 70% EtOH with an ultrasonic bath for 20s. Afterward, they were attached to glass slides with double-sided adhesive carbon tapes and dried at room temperature. Then, each mandible was surrounded by a small metallic ring which was then filled with epoxy resin (Reckli Epoxy WST, RECKLI GmbH, Herne, Germany). After polymerization, lasting for three days at room temperature, glass slides, and tapes were removed. Each sample was subsequently polished with sandpapers of different roughness until the regions of interest in cross-section were on display. Then, the surface was smoothed with aluminum oxide on a polishing machine (Minitech 233/333, PRESI GmbH, Hagen, Germany) and each sample was cleaned by an ultrasonic bath in 70%EtOH, lasting 5 minutes to remove the polishing powder. After mounting the embedded samples on scanning electron microscope (SEM) sample holders, they were sputter-coated with platinum (5 nm layer). The platin was necessary as a reference to check if the EDS measurement was correct (e.g., in some cases, no or very high Pt content was detected; these measurements were excluded from analysis).

Measurements took place at the exocuticle of different mandibular cross-sections (Fig. 1) and were performed with an SEM Zeiss LEO 1525 (One Zeiss Drive, Thornwood, New York, USA) equipped with an Octane Silicon Drift Detector (SDD) (micro analyses system TEAM, EDAX Inc., New Jersey, USA). For each measurement, the same settings were used (i.e. an acceleration voltage of 20 kV, working distance, lens opening, etc.). Before analysis, the detector was calibrated with copper. Overall, 395 small areas (no mapping and no point measurements) were investigated by EDS. The following elements were selected in the software to measure their proportions: H (hydrogen), C (carbon), N (nitrogen), O (oxygen), Pt (platinum), Al (aluminum), Ca (calcium), Cl (chlorine), Cu (copper), Fe (iron), K (potassium), Mg (magnesium), Na (sodium), P (phosphorus), S (sulphur), Si (silicon), and Zn (zinc). Some elements were not discussed here as they are either the elemental basis of chitin and proteins (H, C, N, O), the coating (Pt), or the polishing powder (Al, O). We also performed 10 EDS tests on the epoxy to identify putative pollution due to the mechanical application, embedding, or polishing. We could not detect Si (which is part of the sandpaper), or any other elements, that we further discuss in the resin. Their presence is therefore considered part of the mandible. The peak of P overlaps with one of Pt. Because of this, the software could not discriminate between these two elements and P content could not be reliably determined. Therefore, P and Pt are discussed together (P+Pt). We, however, measured 20 areas of pure epoxy to receive values on their Pt content (mean ± SD; 0.16 ± 0.02 atomic %) to further estimate the proportions of P in the cuticle. We only included measurements where the peaks of the elements were higher than the background noise. After EDS analyses, the samples were used for nanoindentation.

### Material properties

Both cuticular H and E can be measured through nanoindentation (Oliver & Pharr 2004). In nanoindentation experiments, the material surface of a structure is indented with an object of specific geometry (here, a Berkovich indenter tip) and a known force, resulting in material deformation. We used a nanoindenter SA2 (MTS Nano Instruments, Oak Ridge, Tennessee, USA) equipped with a dynamic contact module (DCM) head. To estimate the material H and E we employed the continuous stiffness mode, using indentation-generated force-displacement curves from the loading and unloading phases (Oliver & Pharr 2004). All tests were conducted under normal room conditions (relative humidity 28–30%, temperature 22–24 °C), with each indent and corresponding curve manually controlled. E and H were determined at penetration depths of 600–1000 nm. We received ∼50 values for each site indented, obtained at different indentation depths, which were averaged to receive one H and one E mean value per indent.

Nanoindentation tests were run on the cross-sections of the mandible exocuticle (Fig.1) from the three *F. cunicularia* workers used for EDS analyses (six mandibles were studied). The indentations were applied to the same localities that were tested by EDS analyses, allowing us to relate elemental composition with mechanical properties. After a region of interest was tested by EDS and nanoindentation, the sample was polished and smoothened again until the next target region was on display. In total, 395 sites were tested by EDS and indentation.

### Finite element analysis (FEA)

Generation of mandible volumetric mesh and FEA were conducted in the open-source software FEBio (Maas et al. 2012). We simulated four biting scenarios, namely strike and pressure with the entire masticatory margin or the apical tooth only (Fig.S1). A strike bite consists of a fast-mandibular impact against an object, while a pressure bite simulates the behavior of crushing an object with the mandibles. For strike biting, we applied the load on the apical tooth or the masticatory margin and restricted to zero the nodal displacement in all directions on the mandibular articulations with the head, which maintained the mandible fixed during biting (Fig.S1). For pressure biting, we applied a load on the region of mandibular apodeme origin, representing the insertion of the mandibular closing muscles and its contraction. Also, nodal displacement on the mandibular articulations with the head along with the apical tooth or masticatory margin were restricted to zero in all directions (Fig.S1). A load of 100000 nN was applied in all simulations. We applied the measured E to all pre-defined mandibular regions to compare FEA results with simulations considering a homogeneous E distribution, defined by the value of the mandibular blade. We compared FEA results through color maps and stress intervals (Marcé-Nogué et al. 2017) between E treatments. We considered the von Mises failure criterion (Özkaya et al. 2017) to define a unique stress value for each element.

### Statistical analysis

We applied the intervals method (Marcé-Nogué et al. 2017) to compare the distribution of von Mises non-normalized stress values between each biting simulation with a heterogeneous E distribution and its homogeneous counterpart. Therefore, we compared the proportion of mandibular volume filled by a defined range of stress values at each biting simulation through a Principal Component Analysis (PCA) (Marcé-Nogué et al. 2017). We extracted data on von Mises stress and volume from the elements of each simulation from FEBio (Maas et al. 2012). Then we removed elements representing the 2% higher stress values in each simulation, as these values often represent artificially high-stress values (Marcé-Nogué et al. 2016; 2017). We log-transformed stress values before generating stress intervals to account for variation in the scale of non-normalized von Mises stress values. We defined the upper threshold value to let the 25% higher stress values above the threshold, representing the highest stress interval. For the definition of the ideal number of stress intervals, we generated datasets with different numbers of intervals (5, 10, 15, 25, 50) and performed PCAs. We considered the PC1 and PC2 scores of each dataset in linear regressions with the scores of equivalent PCs of the next interval (e.g., PC1_5__intervals ∼ PC1_10__intervals), and we retrieved the coefficient of determination (R^2^) to analyze the convergence of PC scores. The stop of the R^2^ increase defines the final number of intervals (Marcé-Nogué et al. 2017). Convergence occurred with 15 intervals, so we used this number for the PCAs. We conducted the PCA with the R (R Core Team 2023) packages FactoMineR version 2.4 (Lê et al. 2008) and factoextra version 1.0.7.999 (Kassambara & Mundt 2020).

We tested for differences in element atomic proportion and material properties between the distinct mandibular regions with Kruskal-Wallis tests, applying Dunn tests with Bonferroni corrections to access paired differences between mandibular regions. We also tested for differences in non-normalized stress values of each mandibular region between the E treatments with Mann-Whitney tests, also excluding the 2% higher stress values of each simulation as performed for the intervals method procedure. Statistical analysis were carried out in R (R Core Team 2023) environment, where we employed the package ggstatsplot (Patil 2021) to generate violin plots and perform the Kruskal-Wallis and pair-wise comparison tests.

## Results

### Mandible exocuticle elemental composition

The SRμCT scans of F. cunicularia mandibles showed a disgustingly white band in the mandible cutting edge (Fig.2), suggesting the presence of heavier elements as metals that increase the material density in this region.

**Fig.2.**
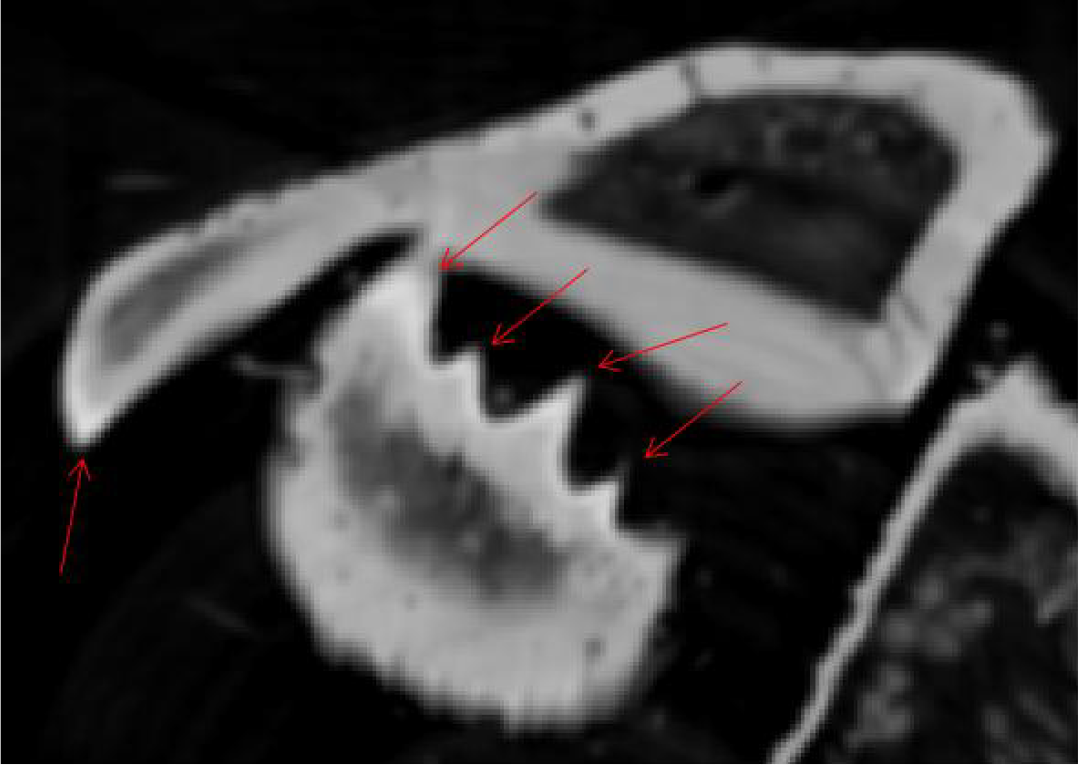
SRμCT scan slice showing the left and right mandibles of a *F. cunicularia* worker highlighting the brighter contour of the masticatory margin (red arrows), which suggest the deposition of materials that increase the density of that region.

According to the EDS results, some atomic elements showed a higher proportion in specific mandibular regions. Notably, Zn, Cu, and P+Pt were found at higher proportions in the mandibular masticatory margin, followed by the DMA, and VMA, showing lower levels in the mandibular blade (Fig.3). Other elements showed differential concentrations at specific mandibular regions (*i.e*. Cl, F, Fe, K, Mn, and Si; Fig.S2), while others were present at similar levels along the entire mandible (*i.e*. Ca, Mg, Na, and S; Fig.S2).

**Fig.3.**
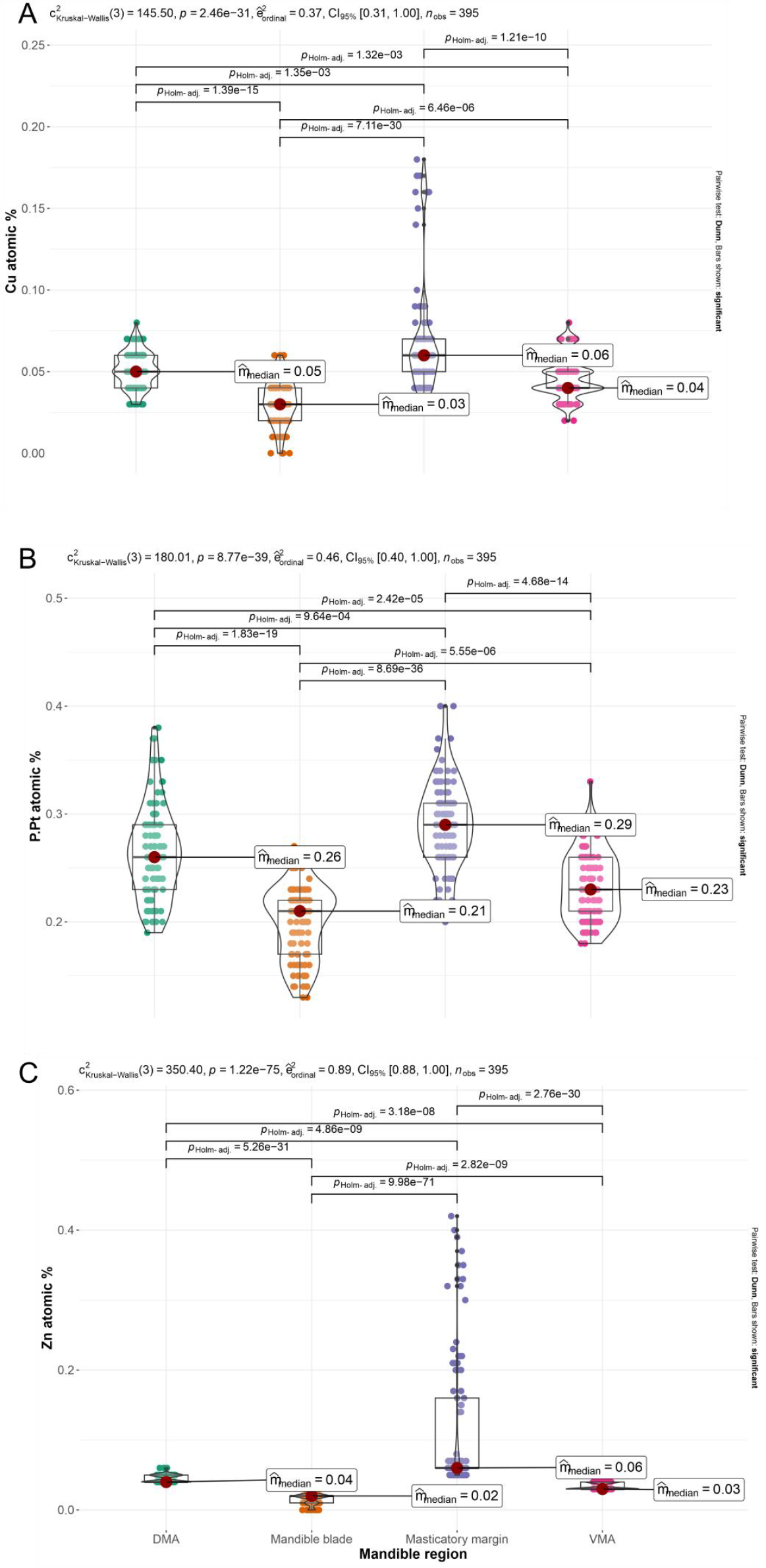
Boxplots depicting the variation in atomic percentage of Cu (A), P+Pt (B), and Zn (C) in distinct regions of *F. cunicularia* worker mandibles, measured with EDS. In the upper left of each graph are depicted the results of Kruskal-Wallis tests for the group difference in atomic % between the mandibular regions, and the horizontal bars connect pairs of mandibular regions that differed in atomic % according to pair-wise Dunn tests (adjusted p-values are shown above the bars). Only significant differences are shown. DMA = Dorsal mandibular articulation; VMA = Ventral mandibular articulation.

### Mechanical properties

Nanoindentation tests showed that the masticatory margin of *F. cunicularia* worker mandibles has higher values of exocuticular H and E (Fig.4), which agrees with the pattern of cuticle density contrast depicted in the SRμCT scans (Fig.2). The mandibular articulations with the head also showed higher exocuticular H and E, with the VMA showing higher values than the DMA of both material properties (Fig.4). Finally, the more delicate mandibular blade showed the lowest levels of exocuticular H and E (Fig.4).

**Fig.4.**
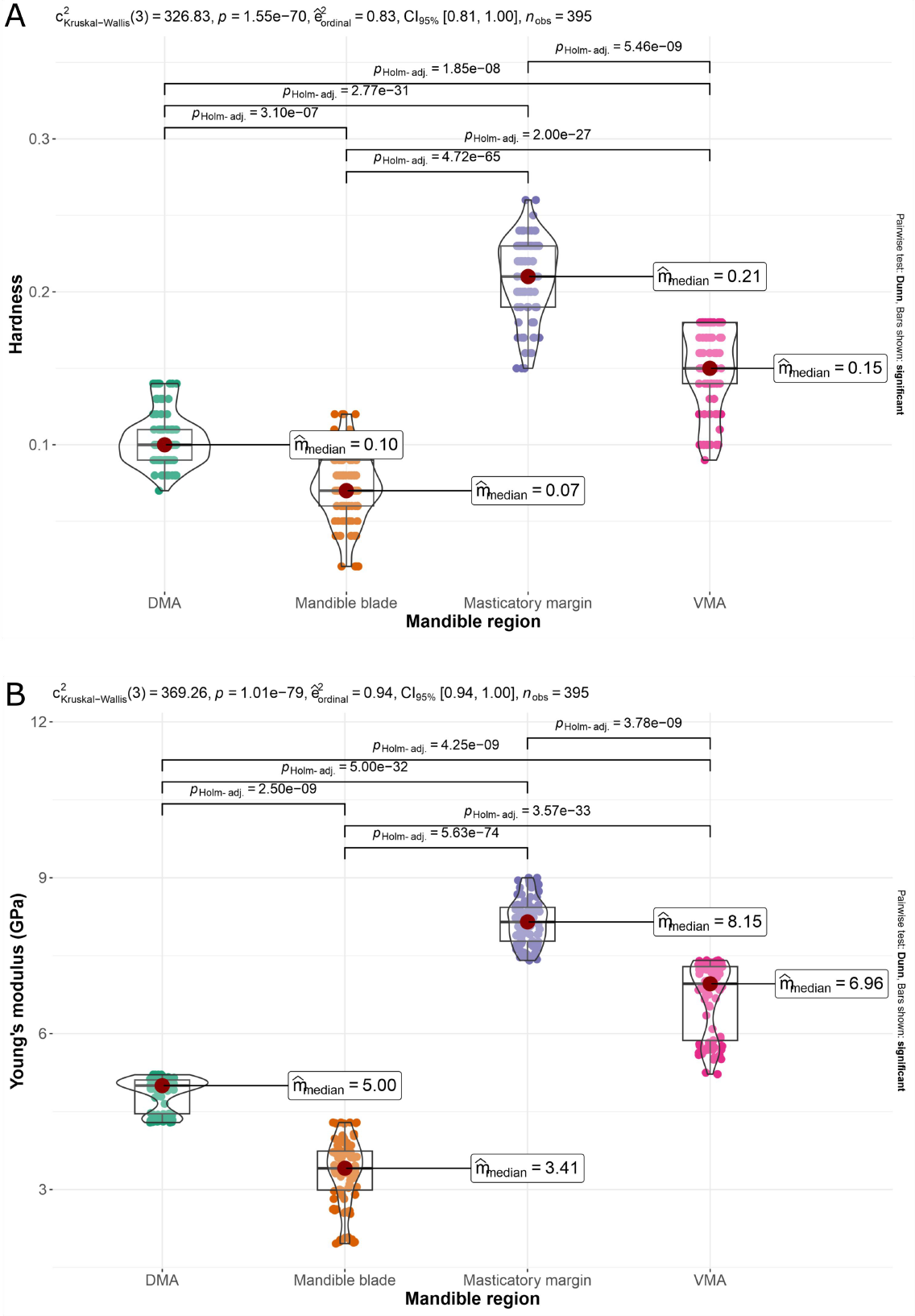
Boxplots depicting the variation in cuticular H (A) and E (B) in distinct mandibular regions of *F. cunicularia* workers, measured with nanoindentation. In the upper left of each graph are depicted the results of Kruskal-Wallis tests for the group difference between the mandibular regions, and the horizontal bars connect pairs of mandibular regions that differed in the respective mechanical properties according to pair-wise Dunn tests (adjusted p-values are shown above the bars). Only significant differences are shown. DMA = Dorsal mandibular articulation; VMA = Ventral mandibular articulation.

### Relationship between parameters

We found relevant correlations between the proportion of some atomic elements and cuticle material properties. For cuticular H, Zn showed the highest correlation (Spearman’s ρ = 0.70). Regarding E, Zn (ρ = 0.77), P+Pt (ρ = 0.54), and Cu (ρ = 0.50) showed the highest correlation values. There were also relevant correlations between atomic elements. The presence of Zn was correlated with P+Pt (ρ = 0.69) and Cu (ρ = 0.64). The occurrence of F correlates negatively with the presence of K (ρ = -0.51) and Mn (ρ = -0.54) and positively with Mg (ρ = 0.62). The distribution of K was correlated with Cl (ρ = 0.50). Finally, the occurrence of S and Ca was also correlated (ρ = 0.67; Fig.5).

**Fig.5.**
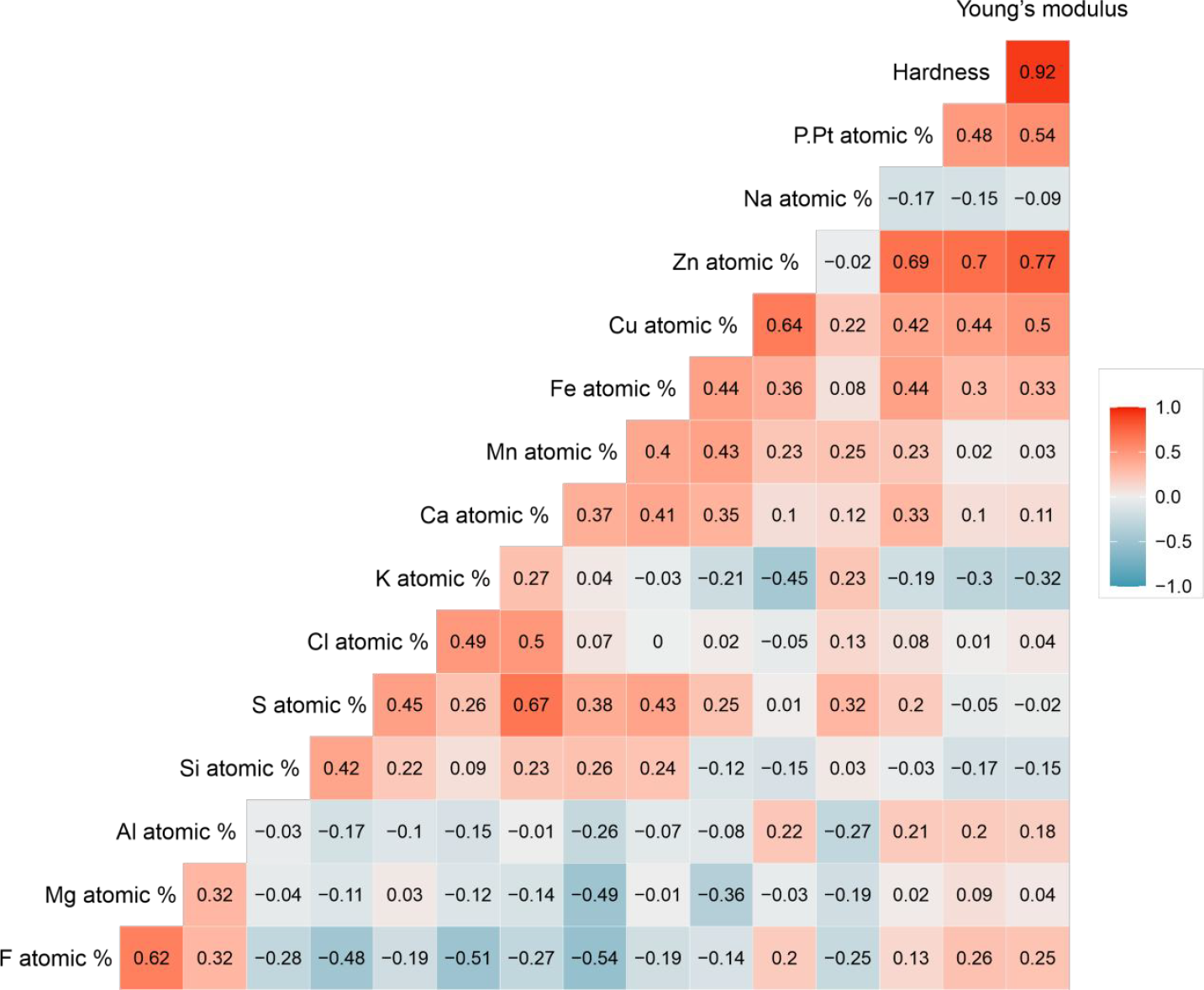
Spearman’s rank correlations among cuticular material properties (H and E) and the proportion of elements for both mandibles of two *F. cunicularia* workers. Bluish and reddish colors represent negative and positive correlations, respectively.

### FEA

The FEA color maps depict subtle but consistent differences in stress patterns between simulations with homogeneous and heterogeneous E (Fig.6). In general, in simulations under a heterogeneous E definition, stresses tended to concentrate at higher levels at the masticatory margin and mandibular articulations, and decrease in the less stiff mandible blade, mainly around the DMA and VMA, along with the masticatory margin (Fig.6). The effect of variation in E on the VMA was more relevant in strike biting (Fig.6e-h). For the masticatory margin, all biting behaviors, except for strike with the entire masticatory margin, showed relevant differences in stress patterns in this region under different E treatments (Fig.6). Regarding the DMA, stresses on the mandibular external face increase when the cuticle was modeled as a homogeneous material, mainly in simulations eploying the entire masticatory margin (Fig.6a-b; e-f).

**Fig.6.**
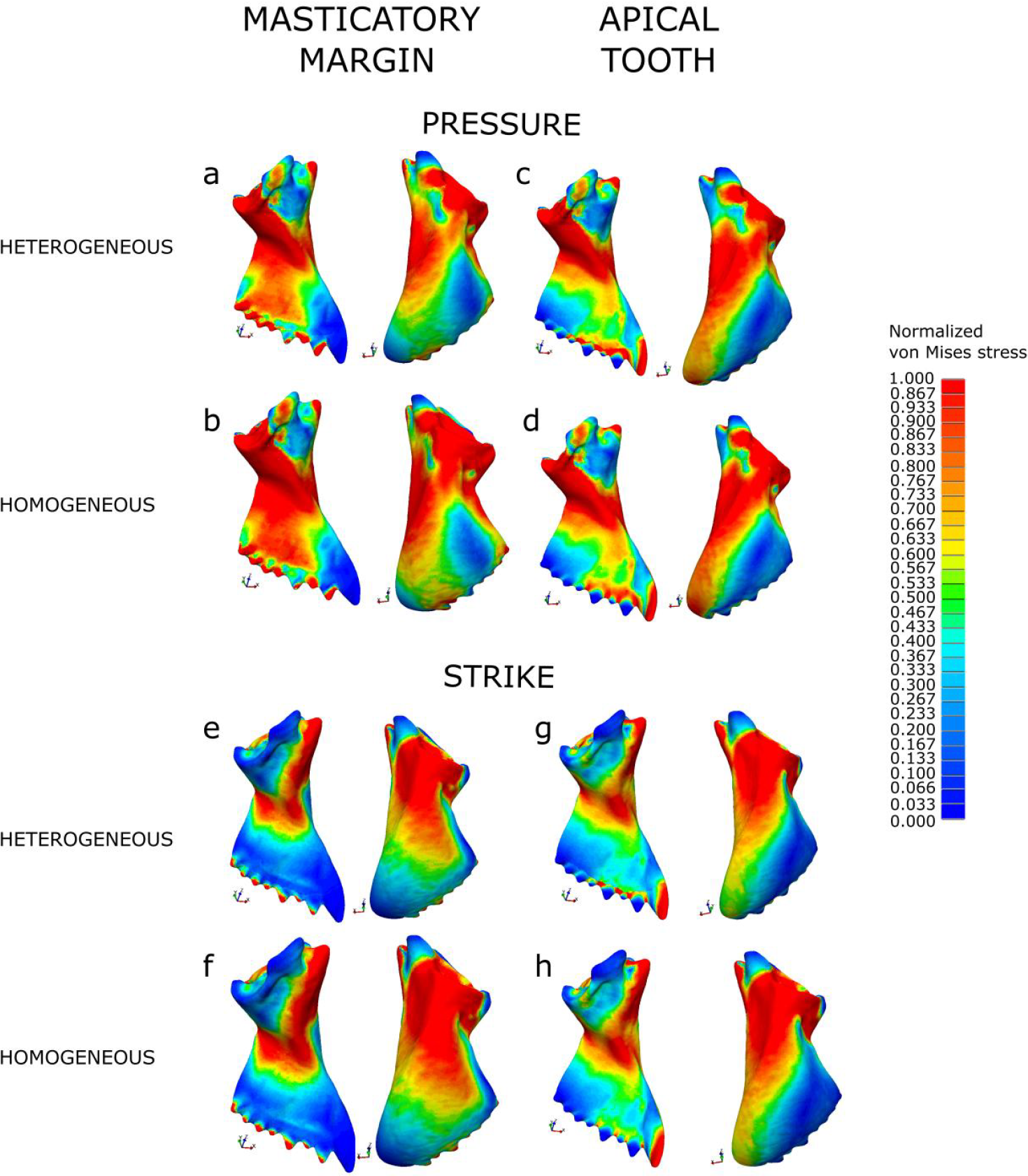
Color maps depicting von Mises stress distribution along the mandibles for all simulations. Stress values were normalized to allow for direct comparisons of stress distribution between simulations. Pressure with the entire masticatory margin (a-b) or the apical tooth only (c-d) under heterogeneous (a and c) or homogeneous (b and d) E; strike with the entire masticatory margin (e-f) or the apical tooth only (g-h) under heterogeneous (e and g) or homogeneous (f and h) E.

When considering the distribution of stress intervals, the first two components of the PCA explained more than 94% of the variance. PC1 was associated with a range of stress intervals from high towards low and intermediate intervals, while PC2 reflects a range of stress intervals from the highest towards the remaining high-stress intervals (Fig.7). The effects of modeling the mandible cuticle as a homogeneous or heterogeneous material were more relevant under strike biting with the masticatory margin, followed by pressure bite in general (Fig.7). Differences between material conditions in strike bite with the masticatory margin were associate with PC1, while for pressure biting the material properties effects were more relevant along PC2 (Fig.7). Irrespective of E treatment, pressure biting was more associated with the lowest stress intervals, and strike biting was more related to the highest stress intervals, mainly when only the apical tooth was employed for biting (Fig.7).

**Fig.7.**
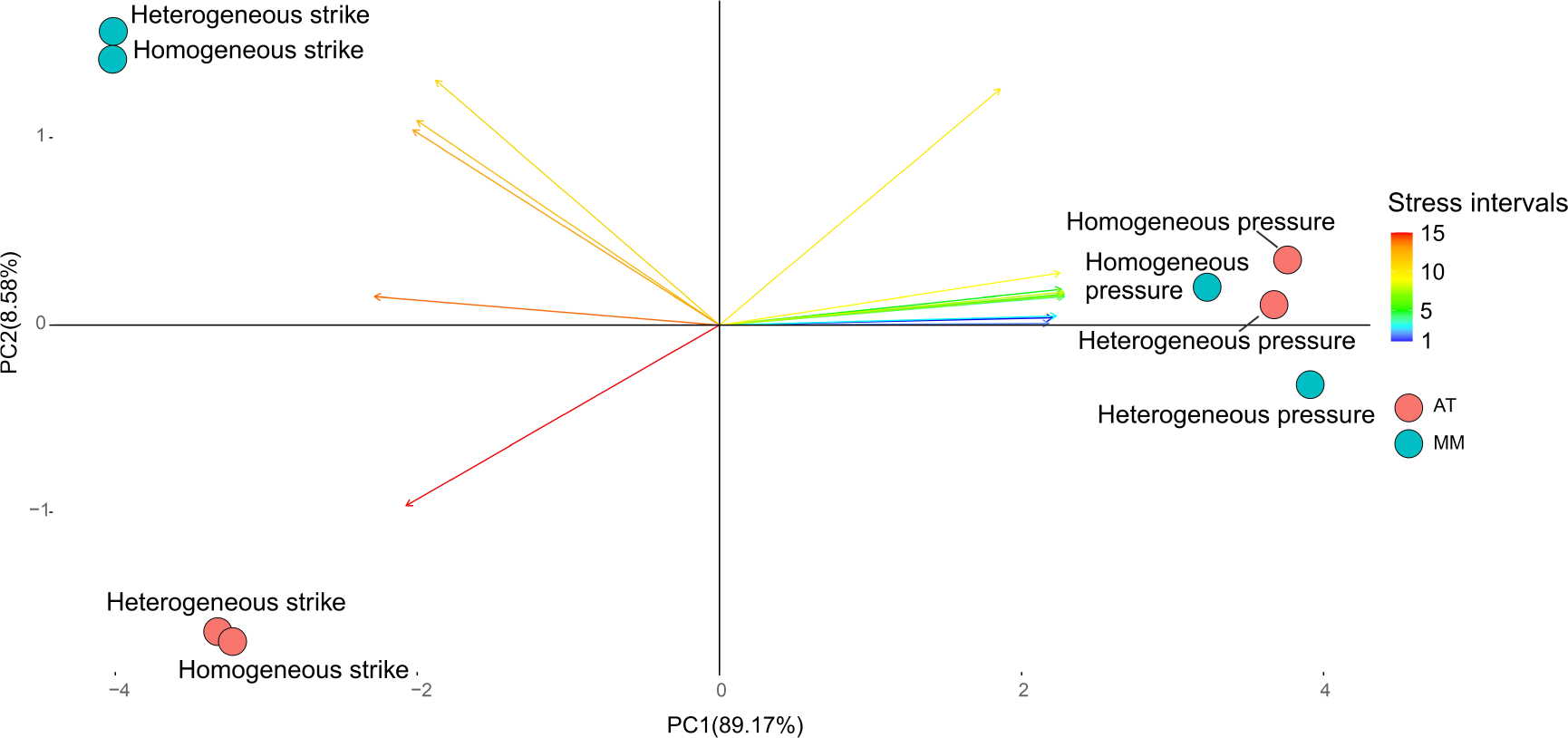
PCA of the first two components depicting the distribution of von Mises stress intervals for all biting simulations. Colored circles represent simulations using the apical tooth only (AT) or the entire masticatory margin (MM). Colored arrows depict distinct stress intervals (from 1 to 15).

Differences in stress patterns were driven by differential stress concentration at each mandibular region between the E treatments. Specifically, there were relevant differences in median stress values in the mandible regions when comparing simulations with homogeneous or heterogeneous E. In strike biting with the masticatory margin, the DMA (*W*_Mann-Whitney_=2.01e+06, p < 0.001) and the masticatory margin (*W*_Mann-Whitney_=3.72e+07, p < 0.001) showed higher stress levels, while the VMA (*W*_Mann-Whitney_=1.80e+05, p = 0.55) and the mandibular blade showed (*W*_Mann-Whitney_=7.19e+09, p = 0.38) no differences when a heterogeneous distribution of E was modeled in comparison to its homogeneous distribution (Fig.S3). A similar pattern was observed in strike biting with the apical tooth only, although in this case, the mandibular blade showed lower stress values under a heterogeneous E distribution than in a homogeneous distribution (Fig.S5). Regarding pressure bite with the entire masticatory margin, the mandible blade again showed lower stress values under a heterogeneous E distribution in comparison to a homogeneous distribution (*W*_Mann-Whitney_=6.70e+09, p < 0.001), and the masticatory margin showed higher stress levels (*W*_Mann-Whitney_=3.69e+07, p < 0.001), as was the case for the VMA (*W*_Mann-Whitney_=4.07e+05, p = 0.02), while for DMA there was no difference between E treatments (*W*_Mann-Whitney_=1.92e+06, p = 0.55) (Fig.S4). The same general pattern was observed in pressure biting with the apical tooth only (Fig.S6).

## Discussion

As expected, our elemental and mechanical characterization of *F. cunicularia* worker mandibles demonstrated that distinct mandibular regions have varying elemental and mechanical attributes, reflecting in mandible responses to bite-loading. The mandibular masticatory margin showed, in general, a higher material density, being brighter than the remaining mandible. This mandibular region was stiffer and harder than the remaining mandible, and showed a higher concentration of Zn, Cu, and P+Pt. When the measured heterogeneity in cuticular E was considered on bite simulations, this region tended to accumulate more stress that otherwise would spread towards the more delicate mandibular blade, similar to snail teeth (Krings et al. 2020).

Interestingly, the mandibular articulations with the head also showed relevant distinctions compared to the mandible blade and masticatory margin. Both DMA and VMA were stiffer and harder and possessed higher levels of Zn, Cu, and P+Pt than the mandible blade, showing lower levels of all those attributes than the masticatory margin. In biting simulations under a heterogeneous E, DMA and VMA also concentrated higher stresses that under a homogeneous E would spread toward the mandible blade. The mandible blade showed the lowest levels of cuticular H and E and concentrated a lower proportion of Zn, Cu, and P+Pt regarding the remaining mandible. When a homogeneous distribution of E was considered in the bite simulations, this mandibular region showed relatively higher stress levels, mainly around the masticatory margin and the mandibular articulations with the head, compared to simulations with a heterogeneous distribution. In general, the stiffer mandibular regions (masticatory margin, VMA, and DMA) tended to concentrate higher stress levels under a heterogeneous E distribution, decreasing the amount of stress spreading towards the mandible blade, when compared to a homogeneous distribution of E.

A higher degree of cuticular H and E along the mandible masticatory margin is commonly observed in ants (Brito et al. 2017; Schofield et al. 2002; 2021), although only a few species were measured so far, mainly leaf-cutter ants. Such higher levels of material properties along the masticatory margin usually follow the concentration of transition metals like Zn (Schofield et al. 2021), which tend to concentrate along the cutting edge or other cutting and piercing tools of arthropods (Polidori et al. 2020; Schofield et al. 2021; Krings & Gorb 2023; Reiter et al. 2023; Krings & Gorb 2023). In leaf cutter ants, the mandible cutting edge is usually stiffer and harder (Schofield et al. 2002; 2021), which could be a mechanism to deal with the abrasion associated with the behavior of cutting leaves, as well as to improve the self-sharpening mechanism of such a structure (Martínez et al. 2020). Worker mandibles wear off during their lifetimes, and older workers with less sharp mandibles tend to change their task rules in the colony to focus on carrying leaves or other activities not related to leaf cutting (Schofield et al. 2011), due to the increased amount of forces needed to cut leaves employing worn mandibles (Püffel et al. 2023b). In *F. cunicularia*, which is not a leaf-cutter ant, the worker masticatory margin is consistently harder and stiffer than the remaining of the mandible, showing also increased levels of Zn, Cu, and P+Pt, followed in those aspects by the mandibular articulations with the head. Distinct from the specialized leaf-cutter ants, *F. cunicularia* has a morphologically monomorphic worker caste, whose foragers exhibit a generalist behavior (Novgorodova 2015). Also, this species usually ranks at a low position in competitive hierarchies, avoiding physical conflict with other ant species (Seifert & Schultz 2009). Our results demonstrate that even a generalist and non-aggressive ant species invest in transition metal accumulation in their worker mandibles, leading to a heterogeneous distribution of material properties with functional relevance for its biting mechanics, suggesting that such patterns may be widely present in other ant lineages.

The evolution of a dicondylic mandibular articulation with the head occurred very anciently in insects, being prevalent in current lineages (Blanke et al. 2015; Blanke 2019), although some reversals to the ancestral condition with a unique point of articulation were suggested (van de Kamp et al. 2022). A dicondylic joint reduces the mandibular movement to a single rotation axis but provides an increased stabilization relevant to generate stronger bites (Gorb & Beutel 2000; Blanke et al. 2015). By being constrained, joint regions generate reaction forces that impose substantial mechanical demands on the associated structures. Although demonstrated that such reaction forces are relevant for insects during bite (Blanke et al. 2017a), there was no attempt so far to characterize the ant mandible cuticle considering the possibility of the mandibular joints being differentially stiffer than other mandibular regions. Accordingly, the functional relevance of modeling material heterogeneity was demonstrated for some organisms and structures (Rajabi et al. 2017; Das et al. 2018; Jafarpour et al. 2020; Li et al. 2020; Matsumura et al. 2020; Krings et al. 2020; Casey et al. 2022), but rarely so regarding bite mechanics. Our results suggest that stiffer regions of the mandible, like the masticatory margin and mandibular joints, can concentrate higher stress levels, reducing the stresses that achieve the more delicate mandibular blade. These stress patterns based on specific material properties’ distributions are important to fulfill the function of cutting and chewing when high stresses are required at the masticatory margin. At the same time, the surrounding cuticle with a lower elasticity modulus acts as a damper to prevent high-stress distribution to the rest of the mandible.

Differences in stress patterns between heterogeneous and homogeneous biting simulations were subtle, demanding a range of FEA results that included color maps, distribution of stress intervals, and differences in mean stress levels among the mandibular regions to be evaluated. However, even little differences in stress concentrations might be decisive for the structure’s proper functioning and simultaneous damage risk minimization. The fact that we were able to demonstrate that the material heterogeneity of the *F. cunicularia* worker mandible has functional relevance suggests that such an approach should be applied to other ant species, especially specialized ants in terms of biting demands, like leaf-cutter and trap-jaw ants, whose bite mechanics was poorly explored besides the relevance of the mandible morphological variation so far (Larabee et al. 2018; Klunk et al. 2021; Wang et al. 2022).

Our results draw attention to the necessity to explore more widely the variation in cuticular elemental and mechanical characteristics of chewing insect mandibles, especially to investigate their functional relevance under bite-loading. Ant workers use their mandibles to perform roughly all non-reproductive tasks they are responsible for, characterizing the multitasking aspect of this working tool (Zhang et al. 2020; Richter & Economo 2023). Although the role of mandibular morphological variation proves to be relevant to bite mechanics (Larabee et al. 2018; Klunk et al. 2021; Püffel et al. 2021; 2023a; Wang et al. 2022), we know little about how the mandible cuticle microstructure can influence its mechanical responses to task-related loading demands.

## Acknowledgments

The authors thank Elke Woelken, from the Department of Electron Microscopy, Institute of Cell and Systems Biology of Animals, Universität Hamburg, for her support at the SEM and Dr. Frank Friedrich, from the Department of Electron Microscopy, Institute of Cell and Systems Biology of Animals, Universität Hamburg, for providing access to the EDS setup to our experiments. This study was financed in part by the Coordenação de Aperfeiçoamento de Pessoal de Nível Superior - Brasil (CAPES) - Finance Code 001, and by the Deutsche Forschungsgemeischaft (DFG) – grant 470833544.

## Authors contribution

C.L.K., M.H., and W.K. conceived the idea of the manuscript; C.L.K., J.U.H., S.N.G., M.H. and W.K. designed the methodology; J.U.H. and W.K. collected the data; C.L.K. and W.K. analyzed the data; C.L.K. and W.K. led the writing of the manuscript. All authors contributed critically to the drafts and gave final approval for publication.

