## Supplementary material for "Mechanical and elemental characterization of ant mandibles: consequences for bite mechanics": Fig.S1

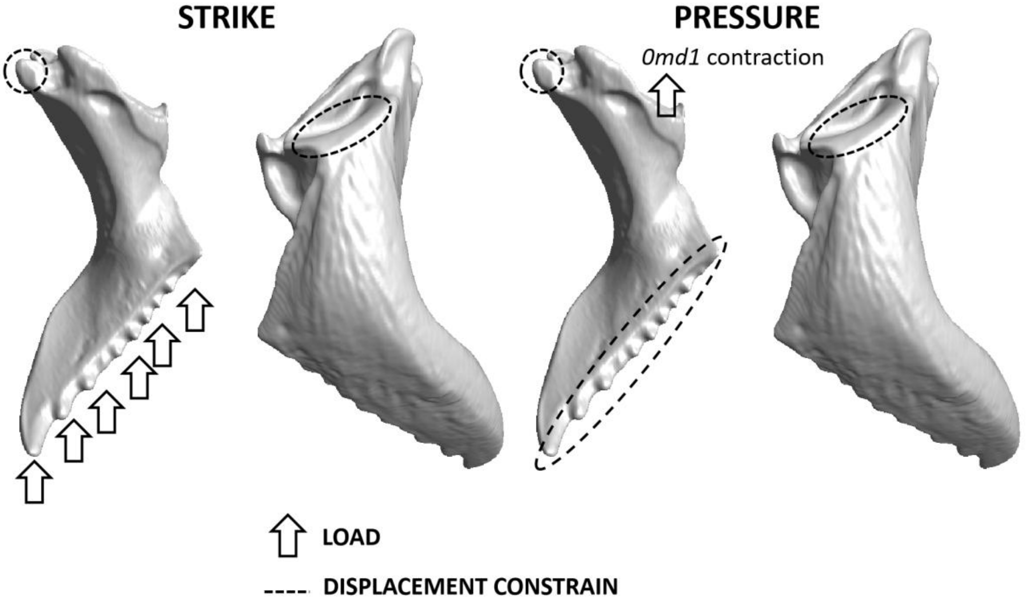


Fig.S1. Diagrams depicting the boundary conditions for each biting scenarios. Highlighted are the conditions for strike and pressure biting with the entire masticatory margin.

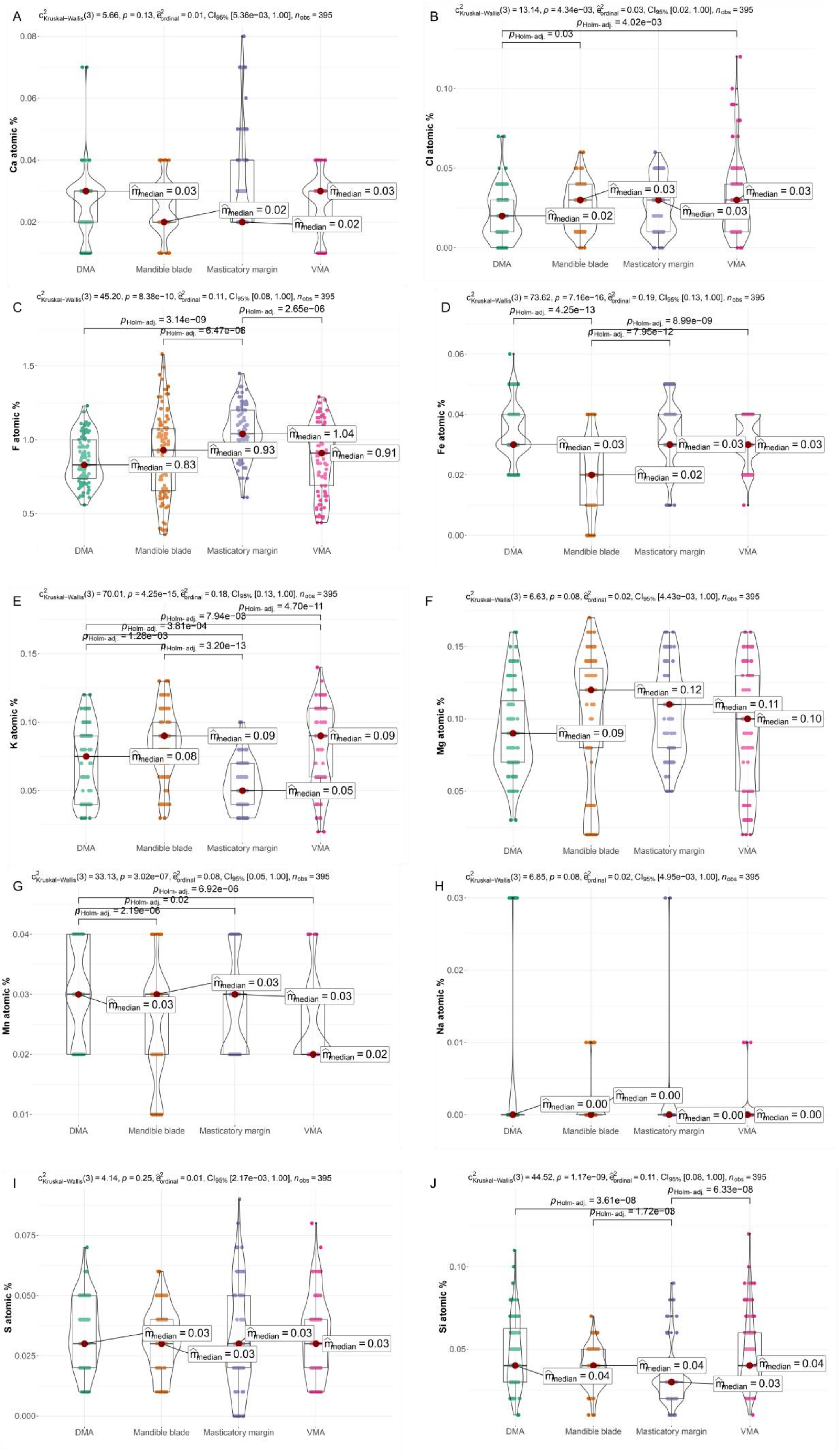

A
c2 Kruskal–Wallis(3) = 5.66, p = 0.13, ε2 ordinal = 0.01, CI95% [5.36e−03, 1.00], nobs = 395
Ca atomic %
m̂median = 0.03
m̂median = 0.02
m̂median = 0.02
m̂median = 0.03
DMA
Mandible blade
Masticatory margin
VMA
B
c2 Kruskal–Wallis(3) = 13.14, p = 4.34e−03, ε2 ordinal = 0.03, CI95% [0.02, 1.00], nobs = 395
pHolm-adj. = 4.02e−03
pHolm-adj. = 0.03
Cl atomic %
m̂median = 0.02
m̂median = 0.03
m̂median = 0.03
m̂median = 0.03
C
c2 Kruskal–Wallis(3) = 45.20, p = 8.38e−10, ε2 ordinal = 0.11, CI95% [0.08, 1.00], nobs = 395
pHolm-adj. = 2.65e−06
pHolm-adj. = 3.14e−09
pHolm-adj. = 6.47e−06
F atomic %
m̂median = 0.83
m̂median = 0.93
m̂median = 1.04
m̂median = 0.91
D
c2 Kruskal–Wallis(3) = 73.62, p = 7.16e−16, ε2 ordinal = 0.19, CI95% [0.13, 1.00], nobs = 395
pHolm-adj. = 4.25e−13
pHolm-adj. = 8.99e−09
pHolm-adj. = 7.95e−12
Fe atomic %
m̂median = 0.03
m̂median = 0.02
m̂median = 0.03
m̂median = 0.03
E
c2 Kruskal–Wallis(3) = 70.01, p = 4.25e−15, ε2 ordinal = 0.18, CI95% [0.13, 1.00], nobs = 395
pHolm-adj. = 4.70e−11
pHolm-adj. = 7.94e−08
pHolm-adj. = 3.81e−04
pHolm-adj. = 1.28e−03
pHolm-adj. = 3.20e−13
K atomic %
m̂median = 0.08
m̂median = 0.09
m̂median = 0.05
m̂median = 0.09
F
c2 Kruskal–Wallis(3) = 6.63, p = 0.08, ε2 ordinal = 0.02, CI95% [4.43e−03, 1.00], nobs = 395
Mg atomic %
m̂median = 0.09
m̂median = 0.12
m̂median = 0.11
m̂median = 0.10
G
c2 Kruskal–Wallis(3) = 33.13, p = 3.02e−07, ε2 ordinal = 0.08, CI95% [0.05, 1.00], nobs = 395
pHolm-adj. = 6.92e−06
pHolm-adj. = 0.02
pHolm-adj. = 2.19e−06
Mn atomic %
m̂median = 0.03
m̂median = 0.03
m̂median = 0.03
m̂median = 0.02
H
c2 Kruskal–Wallis(3) = 6.85, p = 0.08, ε2 ordinal = 0.02, CI95% [4.95e−03, 1.00], nobs = 395
Na atomic %
m̂median = 0.00
m̂median = 0.00
m̂median = 0.00
m̂median = 0.00
I
c2 Kruskal–Wallis(3) = 4.14, p = 0.25, ε2 ordinal = 0.01, CI95% [2.17e−03, 1.00], nobs = 395
S atomic %
m̂median = 0.03
m̂median = 0.03
m̂median = 0.03
m̂median = 0.03
J
c2 Kruskal–Wallis(3) = 44.52, p = 1.17e−09, ε2 ordinal = 0.11, CI95% [0.08, 1.00], nobs = 395
pHolm-adj. = 6.33e−08
pHolm-adj. = 3.61e−08
pHolm-adj. = 1.72e−08
Si atomic %
m̂median = 0.04
m̂median = 0.04
m̂median = 0.03
m̂median = 0.04

Fig.S2. Boxplots depicting the variation in atomic percentage of Ca (A), Cl (B), F (C), Fe (D), K (E), Mg (F), Mn (G), Na (H), S (I), and Si (J) in distinct regions of *F. cunicularia* worker mandibles, measured with EDX. In the upper left of each graph are depicted the results of Kruskal-Wallis tests for the group difference in atomic % between the mandibular regions, and the horizontal bars connect pairs of mandibular regions that differed in atomic % according to pair-wise Dunn tests (adjusted p-values are shown above the bars). Only significant differences are shown. DMA = Dorsal mandibular articulation; VMA = Ventral mandibular articulation.

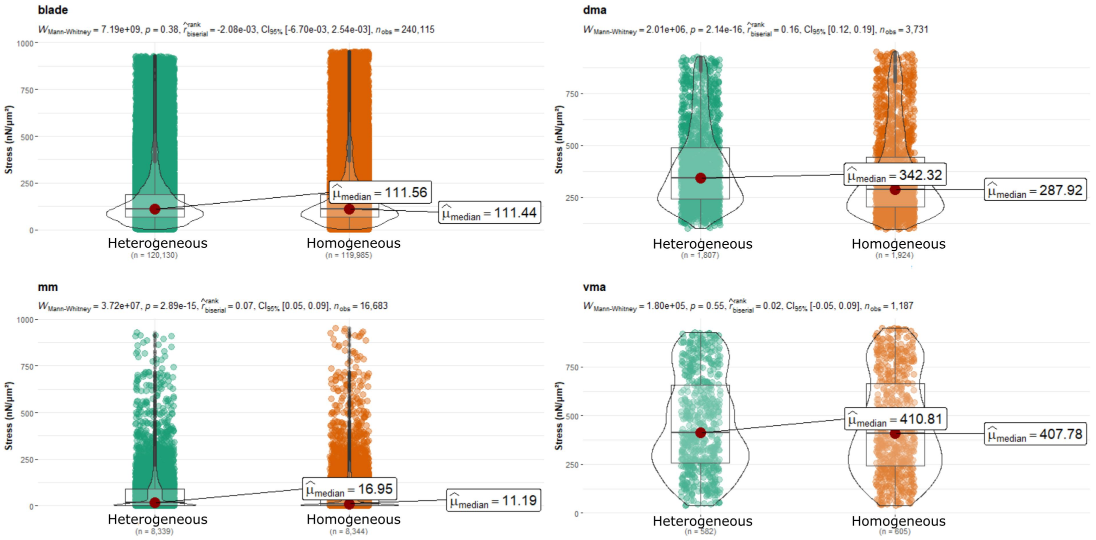


Fig.S3 Violin plots depicting the distribution of non-normalized von Mises stress values in each mandibular region of a mandible with a heterogeneous versus homogeneous E distribution (see main text for details) submitted to strike with the entire masticatory margin. Median stress values are represented by the red dots and the accompanying box. Differences in stress values were accessed with a Mann-Whitney test, whose results are depicted in the upper left of each graph.

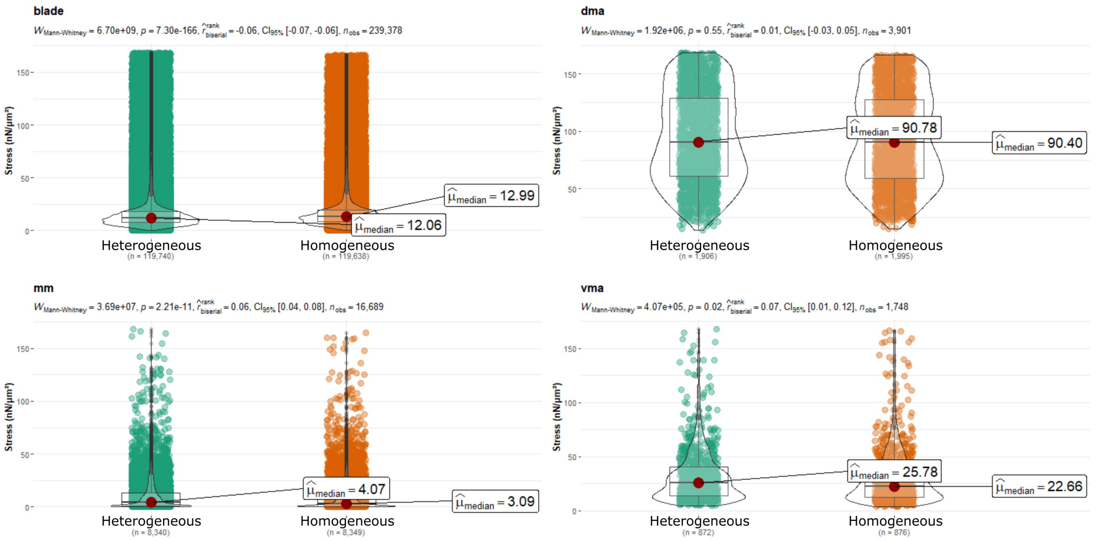


Fig.S4 Violin plots depicting the distribution of non-normalized von Mises stress values in each mandibular region of a mandible with a heterogeneous versus homogeneous E distribution (see main text for details) submitted to pressure with the entire masticatory margin. Median stress values are represented by the red dots and the accompanying box. Differences in stress values were accessed with a Mann-Whitney test, whose results are depicted in the upper left of each graph.

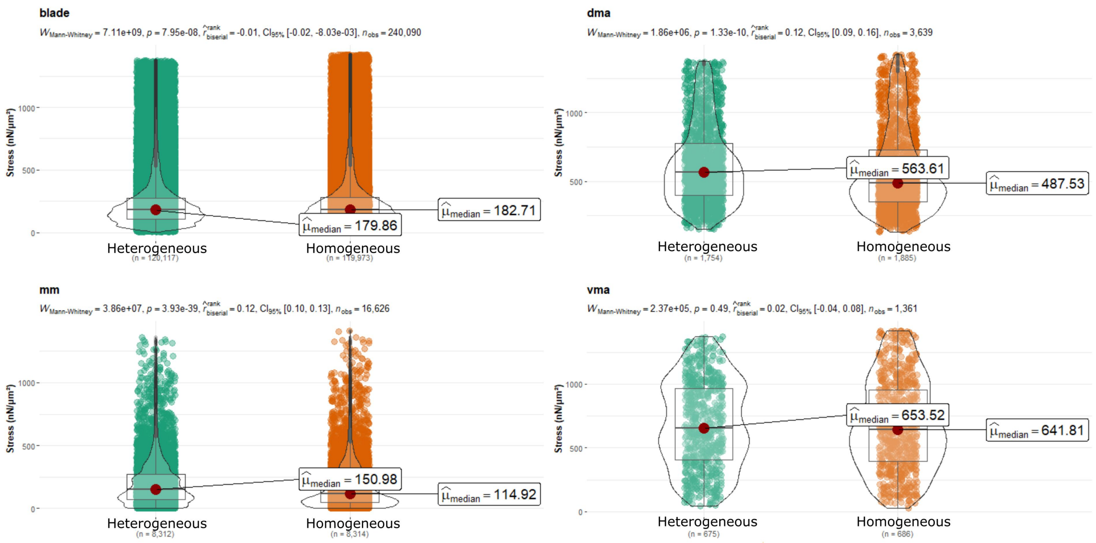


Fig.S5 Violin plots depicting the distribution of non-normalized von Mises stress values in each mandibular region of a mandible with a heterogeneous versus homogeneous E distribution (see main text for details) submitted to strike with the apical tooth. Median stress values are represented by the red dots and the accompanying box. Differences in stress values were accessed with a Mann-Whitney test, whose results are depicted in the upper left of each graph.

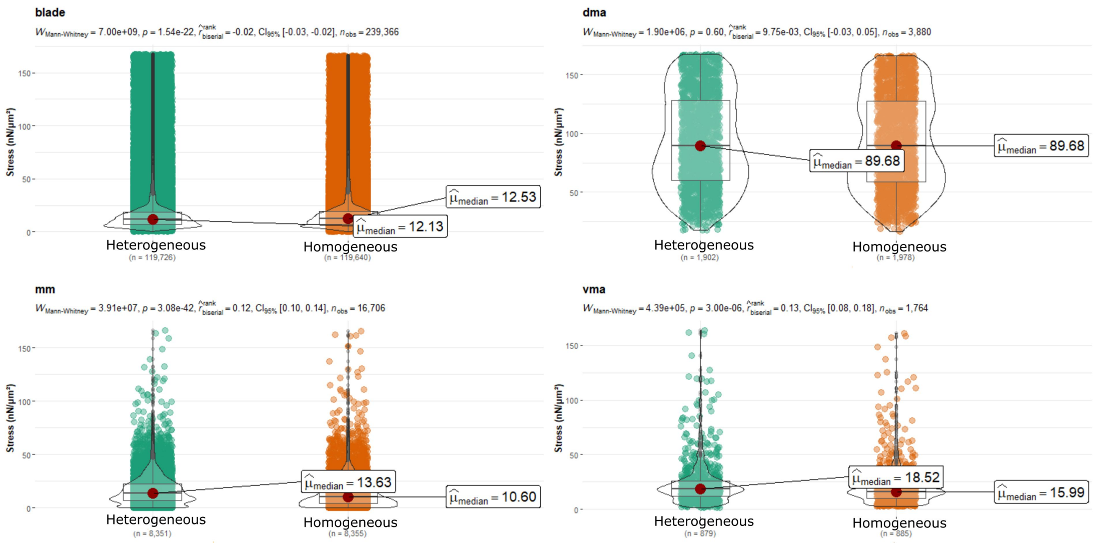


Fig.S6 Violin plots depicting the distribution of non-normalized von Mises stress values in each mandibular region of a mandible with a heterogeneous versus homogeneous E distribution (see main text for details) submitted to pressure with the apical tooth. Median stress values are represented by the red dots and the accompanying box. Differences in stress values were accessed with a Mann-Whitney test, whose results are depicted in the upper left of each graph.
